# Dissociating the functional role of the para-hippocampal and the parietal cortex in human multi-step reinforcement learning

**DOI:** 10.1101/2025.04.15.648746

**Authors:** Fabien Cerrotti, Alexandre Salvador, Sabrine Hamroun, Maël Lebreton, Stefano Palminteri

**Affiliations:** Département d’Études Cognitives, École Normale Supérieure, Université de Recherche Paris Sciences et Lettres, Paris, France; Laboratoire de Neurosciences Cognitives Computationnelles, Institut National de la Santé et Recherche Médicale, Paris, France; CNRS PjSE UMR 8545 & Paris School of Economics, Paris, France; Swiss Center for Affective Sciences, Faculty of Psychology and Educational Sciences, University of Geneva, Geneva, Switzerland

**Keywords:** reinforcement learning, model-based, planning, parietal cortex, temporal lobe, reward system

## Abstract

Many real-life decisions involve a tension between short-term and long-term outcomes, which requires forward-looking abilities. In reinforcement learning, this tension arises at the initial stage of multi-step learning and decision tasks, where forward-looking decisions collect smaller immediate rewards but govern the transition to more advantageous second-stage states. Here, we investigated the neural mechanisms underlying such forward-looking decisions in a cohort of healthy participants undergoing fMRI scanning (N=28). Behavioral results confirmed that participants were able to learn concurrently reward values and state-transition probabilities. By contrasting BOLD signal elicited by first-stage versus second-stage at the time of decision, we isolated a brain network, with central nodes in the bilateral parahippocampal cortex (BA37), whose activity correlate with forward-looking choices. BOLD activity in another network, including the bilateral parietal cortex, correlated with structure learning signal at the time of outcome processing. Our results shed new light on the neural bases of model-based reinforcement learning by suggesting a specific role of the parahippocampal cortex in forward planning and the parietal cortex in learning the Markovian structure of the task.

## Introduction

Complex, ecological environment, which naturally features successions of decisions, actions and consequences, can generally be formalized as Markovian spaces, i.e. network of states in which decisions between available actions define both the probability of transitioning to adjacent states and of collecting state-specific outcomes (Sutton & Barto, 1999). Therefore, navigating such environments involves balancing short-term (present state) and long-term (future states) outcomes, a challenge that is central to animal, human and artificial decision-making (Kahneman, 2011; Mattar & Lengyel, 2022; Miller & Venditto, 2021). Reinforcement learning (RL) consists in a collection of algorithms designed to solve this kind of problems and, in some specifications (model-based RL), assumes that agents build a cognitive model (or ‘map’) of their environment, which represents the probabilistic relationships between different states and between states and actions (Dayan & Daw, 2008; Tolman, Edward Chace, 1951). This allows them to evaluate actions prospectively, integrating immediate results with future rewards obtained after transitioning to new states, and promotes forward-looking decisions (Daw et al., 2005; Gläscher et al., 2010).

The neuro-computational underpinnings of model-based decision-making and learning have been actively investigated using multi-step tasks, where outcomes are delivered after decision-trees involving two or more choices (Daw et al., 2011; Huang et al., 2020; Tanaka et al., 2016). Studies using this kind of tasks originally demonstrated that the ventral striatum is critically involved in processing reward signals, regardless of how these signals are generated (Daw et al., 2011), and that the parietal and the dorsolateral prefrontal cortices are more specifically involved in computing the state prediction errors inherent to model-based learning algorithms (Gläscher et al., 2010). A second wave of studies then highlighted the role of the medial temporal lobe, including the hippocampal and parahippocampal cortex, in model-based decision-making (Bornstein & Daw, 2013; Doll et al., 2015; Kahnt & Tobler, 2016; Wang et al., 2020). Most recently, this literature has been somewhat supplanted by a flourishing of studies dissecting the formation of predictive, cognitive maps in the hippocampus (Garvert et al., 2017; Stachenfeld et al., 2017; Vaidya & Badre, 2022; Whittington et al., 2022; Wikenheiser & Schoenbaum, 2016), a cognitive process that profoundly generalizes the strategies underlying model-based learning, planning and decision-making. These studies have generally leveraged sophisticated experimental paradigms derived from spatial navigation task, which involve more complex computations and representations than the original two-step task (Garvert et al., 2017; Stachenfeld et al., 2017; Whittington et al., 2022; Wikenheiser & Schoenbaum, 2016).

As a consequence, despite these outstanding progresses, the specific roles of the dorsoparietal system and medial temporal structures (including the hippocampal and peri-hippocampal cortices) in multi-step decision-making remain paradoxically unclear. Notably, while the fact that the dorsoparietal system signals the state prediction errors essential to structure learning appears quite consensual, the precise role of the medial temporal lobe (MTL) is less well-defined. Specifically, it is uncertain whether this region is primarily involved in learning or in exploiting the cognitive map for planning and decision-making. On the one hand, influential studies supporting the key role of MTL in the representation and imagination of options for future-oriented choices would suggest that MTL activity is critically at the initial deliberation of a multi-step decision problem, to address the tension between the forward-looking option and the immediate reward (Buckner, 2010; Comrie et al., 2022; Hassabis et al., 2007; Lebreton et al., 2013; Peters & Büchel, 2010). On the other hand, the growing body of literature linking hippocampal activity to generative replay, predictive representations, and credit assignment would suggest that MTL activity is specific to the outcome of the final stage of the same problem (Liu et al., 2021; Stachenfeld et al., 2017).

Critically, previous studies could seldom dissociate the learning from the planning and decision-making aspect of multi-step decision problems, because they mostly focused on contexts where the task structure was either explicitly explained or trained before exposure to rewards (Bornstein & Daw, 2013; Daw et al., 2005; Doll et al., 2015; Gläscher et al., 2010; Hampton et al., 2006). This leaves open the question of how the brain represents and arbitrates among concurrent decision and learning signals in more ecological scenarios where it must simultaneously learn both reward values and task structure.

To address these questions, we employed a novel protocol of the two-step task designed to elucidate the specific roles of different brain regions in multi-step RL. In this task, the optimal policy requires forgoing short-term rewards at the first step to obtain a greater reward at the second step (Daw et al., 2011; Feher da Silva & Hare, 2020; Kool et al., 2016). We included a post-learning assessment to verify that subjects learned the task structure explicitly. Subjects underwent fMRI scanning during the learning task, allowing us to identify brain activations associated with structure learning and utilization through simple contrasts.

Our analyses revealed distinct roles for different brain regions in multi-step RL. We observed that the medial temporal cortex (and specifically the parahippocampal cortex) and dorsoparietal areas are recruited at different stages of the task. Additionally, we found classical reward-related activations in the ventral striatum and ventral prefrontal cortex, modulated by task position, consistent with the notion that rewards are more ambiguously defined in the first step where there is a tension between long-term and short-term outcomes. The present study aims to elucidate the functional roles of the parahippocampal and parietal cortex in human multi-step reinforcement learning, shedding new light on the neural mechanisms underlying forward planning and structure learning.

## Results

### Behavioral design and results

The study used a two-step learning decision task (**Fig. 1A**), which was inspired by previous research (Daw et al., 2011; Gläscher et al., 2010). In each trial, participants were presented with two options represented by abstract symbols (Agathodaimon alphabet). Conditional on being chosen, both options probabilistically determined the transition to one of two second-step states, each represented by distinct colors (**Fig. 1A**). In this second-step, participants were presented with a choice between two new symbols. The number of point (rewards) they collected at each trial was dependent on the combination of chosen options (**Fig. 1B**; see **Table 1** for more details about the contingencies): Options with associated with smaller expected rewards in the first step granted a higher probability of transitioning to a “reward-rich” state in the second step. Conversely, options associated with higher expected rewards in the first step granted higher probability of transitioning to a “reward-poor” state in the second step. Contrary to previous studies, where state-transition probabilities were learned or given through instructions before the introduction of rewards and henceforth of value learning, our experimental setup enables for the evaluation of both structure (transitions) and value learning simultaneously (Daw et al., 2011; Gläscher et al., 2010). In our study, participants performed the task over three sessions and were not given explicit indications about the structure of the two-step task. Also, the presence of rewards in the first step allowing assessing whether the simultaneous learning of task rewards and structure is possible. Note that in the subsequent we will present the results averaged across the behavioral pilot (n=31) and fMRI experiment (n=26), but see **Fig. S1** for the results presented individually.

**Table 1.** Task parametric design of the task. Four groups of participants performed this task inside or outside the MRI. Transition probabilities (*T*(*R*)), rewards probabilities (*P*(*T*)) and rewards (*R*) differed and were counterbalanced between groups and across the three sessions.

| Group | Num. participants | Trials per session | Session 1 |  |  | Session 2 |  |  | Session 3 |  |  |
| --- | --- | --- | --- | --- | --- | --- | --- | --- | --- | --- | --- |
| | | | $P(R)$ | $P(T)$ | $R$ | $P(R)$ | $P(T)$ | $R$ | $P(R)$ | $P(T)$ | $R$ |
| Outside MRI | 15 | 80 | 0.8 | 0.7/1 | $\frac{0}{100} \mid \frac{40}{10}$ | 0.8 | 0.7/1 | $\frac{0}{100} \mid \frac{40}{4}$ | 0.8 | 0.7/1 | $\frac{0}{100} \mid \frac{40}{10}$ |
| Outside MRI | 15 | 80 | 0.8 | 0.8/1 | $\frac{0}{100} \mid \frac{40}{4}$ | 0.8 | 0.8/1 | $\frac{0}{100} \mid \frac{40}{10}$ | 0.9 | 0.9 | $\frac{0}{100} \mid \frac{30}{10}$ |
| Inside MRI | 14 | 80 | 0.8 | 0.8/1 | $\frac{0}{80} \mid \frac{50}{15}$ | 0.8/1 | 0.8 | $\frac{0}{120} \mid \frac{40}{25}$ | 0.8 | 0.8 | $\frac{0}{100} \mid \frac{30}{10}$ |
| Inside MRI | 14 | 80 | 0.8/1 | 0.8 | $\frac{25}{120} \mid \frac{40}{15}$ | 0.8 | 0.8/1 | $\frac{25}{120} \mid \frac{40}{15}$ | 0.8 | 0.8 | $\frac{0}{100} \mid \frac{30}{10}$ |

To evaluate participants’ learning, we first analyzed their long-term choice rate for first step choices in each session (**Fig. 1E**). A long-term choice rate was defined as the frequency with which participant selected the first-step option associated with a lower probability of high reward, yet with a higher probability of transitioning to a reward-rich state at the second step. We observed a significant effect of sessions on the long-term choice rate (*χ*^2^(2) = 13.8, *P* = .001), characterized by a preference for the long-term option which evolved from non-significant in the first two sessions (first: *W*(57) = 493, *P* = .998; second: *W*(57) = 735, *P* = .827) to significant in the third session (*W*(57) = 995, *P* = .030). This suggests that participants gradually switched from a short-term reward maximization to a forward-looking strategy during the task. Crucially, participants displayed a fair level of choice accuracy in the second step choice, indicating that they learned the values of options in both terminal states. Indeed, the correct choice rate was way above chance in the both the “poor” and “rich” (poor *W*(57) = 1706, *P* < .001, *M* = 74.12%; rich *W*(57) = 1711, *P* < .001, *M* = 85.45%; difference: (*t*(57) = 5.84, *P* < .001, *M* = 11.33%; **Fig. 1F**).

As novel features, our experimental design also included two post-learning assessments, allowing to evaluate participants’ learning of the option values and the task structure (**Fig. 1C**). More specifically, in a preference assessment, participants were asked which stimulus they believed to be the more advantageous to them among those displayed on the screen (all possible combinations were displayed). In addition, in a transition assessment task, participants were asked which was the most probable state visited in the second step after choosing a given symbol of the first step (**Fig. 1D**). Note that post-learning assessments were run only on the stimuli of the last session. The evaluation of choice ratios in the post-learning preference assessment reveals a preference for symbols with high rewards in rich states over symbols with high rewards in poor states (**Fig. 1G**; *W*(57) = 5037, *P* < .001), again evidencing those participants learned the task structure. It is also noteworthy that the choice rate for the poor state’s high-value option (P-H) was higher than that of the rich state’s low-value option (R-L; *W*(57) = 2448, *P* < .001), despite an objectively lower expected value – a signature consistent with context-dependent learning (Palminteri & Lebreton, 2021). The correct choice rate for the transitions to both the reward-poor states and to reward rich ones was similar and significantly above chance (**Fig. 1H**; poor: *W*(57) = 950, *P* < .001; rich: *W*(57) = 1058, *P* < .001; difference: *W*(57) = 555, *P* = .320, with a small effect size, *r* = .173), again evidencing that participants learned the task structure.

In conclusion, in our experiment, participants progressively learned to make long-term decisions, such that forward-looking strategy peaked in in the last session of the experiment (**Fig.1 E**). Furthermore, they formed −and were able to retrieve− an explicit representation of the structure of the task, as suggested by the transition assessment results evaluated after this last session (**Fig. 1H**).

### Neural correlates of learning and using a task structure

In order to assess the neural correlates of the cognitive processes engaged in our multi-step decision-making task, we switched to the analysis of the BOLD signal. We designed a first general linear model (GLM1) in which we characterized each of the two steps as two trial events, modelled as stick functions centered around the onsets of the choices and the outcomes (**Fig. 2A**). Choice onsets were parametrically modulated by the reaction times and outcome onsets by the obtained reward (**Fig. 1S**). Each parametric modulator was z-scored to ensure the commensurability of the regression coefficients (Lebreton et al., 2019).

We first compared the activations elicited at the choice onsets of the first step versus the second step. Critically, these two events are rigorously matched on motor and visual characteristic, but forward-looking computations and planning only occur in the first step choices. This contrast revealed a significant positive activation in the bilateral parahippocampal cortex (PHC) (familywise error correction, *P*_*FWE*_ < .05; **Fig. 2B**). This indicates that the PHC is more activated during first step, compared to second step, choices. The opposite contrast revealed significant activations in a prefrontal network involving the dorsal anterior cingulate cortex (dACC) and orbitofrontal cortex (OFC), which were more activated during second step choices (*P*_*FWE*_ < .05; **Fig. 2B**).

**Figure 1.**
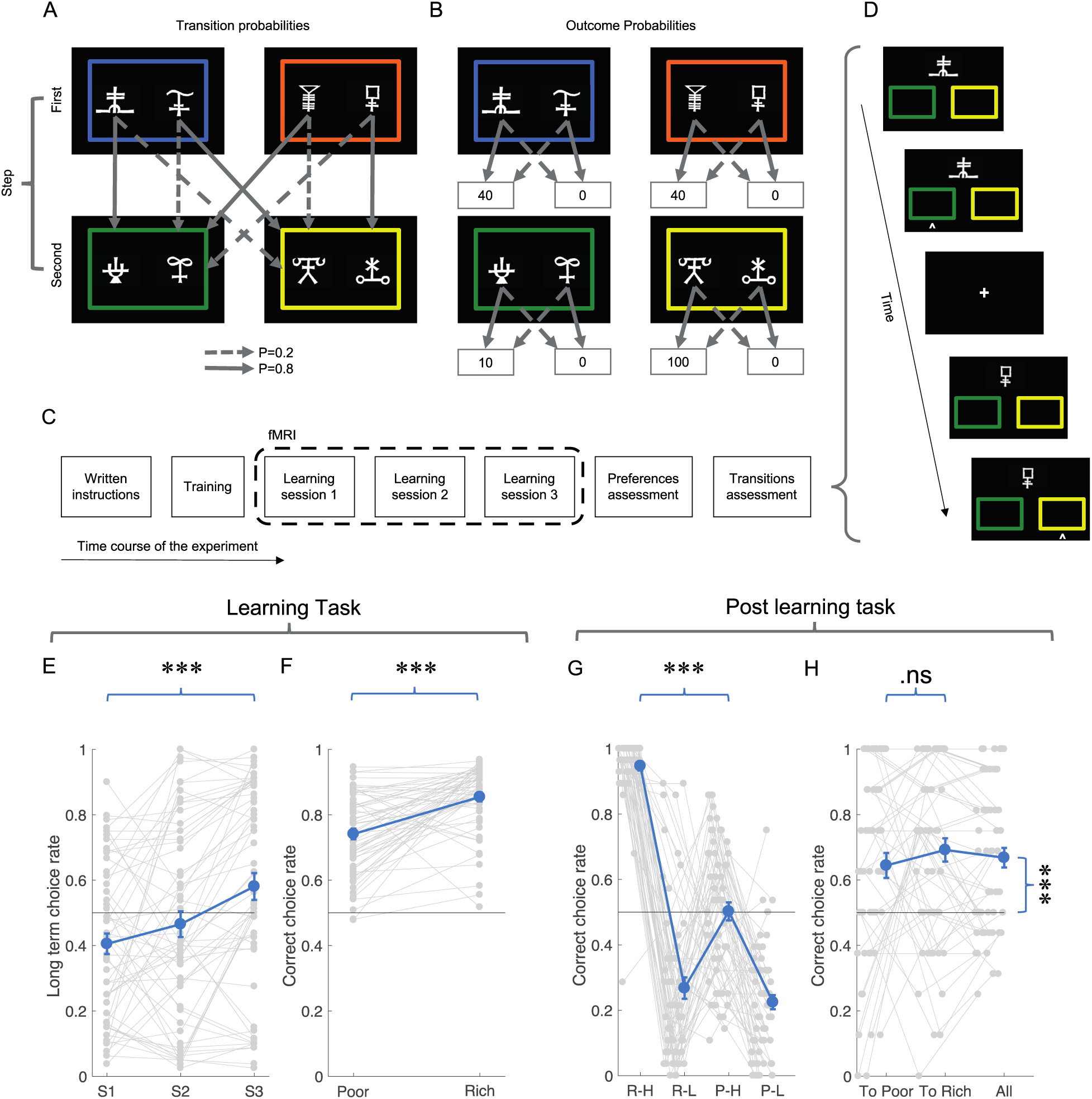
Task design and behavioral results. (**A**) Illustration of the transition probabilities from the first to the second step, depending on the option selected in the first step. (**B**) Illustration of reward probabilities obtained, depending of the choice on the option selected in the first and second steps. (**C**) Task structure: the task involves training, three fMRI sessions, a post-learning preference assessment, and a post-learning transition assessment. (**D**) Trial structure for the post-learning transition assessment: the participant is presented with a symbol from the first step and instructed to estimate whether it leads to the poor state (green) or the rich state (yellow). (**E**) The graph represents the long-term choice rate in the learning task per sessions. (**F**) The graph represents the rate of correct choices based on the state in the second step (Poor or Rich). (**G**) The graph displays the rate at which participants accurately selected the first step option that led to either the poor or rich state in the second step. (**H**) The graph displays the accuracy rate at the second step based on the state (poor/rich) and outcome value (low/high). The gray lines and dots represent the performance of each participant, while the mean is shown in blue. Error bars indicate the standard error of the mean. Significant differences are denoted by asterisks: * for *P* < .05, ** for *P* < .01, and *** for *P* < .001.

We then compared the activity elicited at the outcome onsets of the first step versus the second step (**Fig. 2A**). These events are rigorously matched in basic visual properties (e.g., luminance), but the feedback obtained in the first step is uniquely multidimensional, containing information about both the obtained outcome and the transition the next state. Therefore, while the outcome of the second step only characterizes the learning of option values, the outcome of the first step characterizes the learning of both option values and option-state transitions. The contrast revealed a large cluster of activations encompassing part of the occipital lobe and the parietal cortex (*P*_*FWE*_ < .05; **Fig. 2C**), which were more activated during outcome processing in the fist, compared to the second, step. Additionally, the opposite contrast revealed significant clusters in the insula and medial prefrontal cortex (mPFC), which were more activated during second step outcomes (*P*_*FWE*_ < .05; **Fig. 2C**).

In summary, our simple contrasts revealed distinct neural networks involved in multi-step learning and decision making. More specifically, first step decisions, which mobilize forward looking computations, are associated with activation of the parahippocampal cortex, whereas first step outcomes, which sustain learning of both option values and state transitions, elicit specific activations in a large posterior network encompassing the occipital lobe and the parietal cortices.

**Figure 1.**
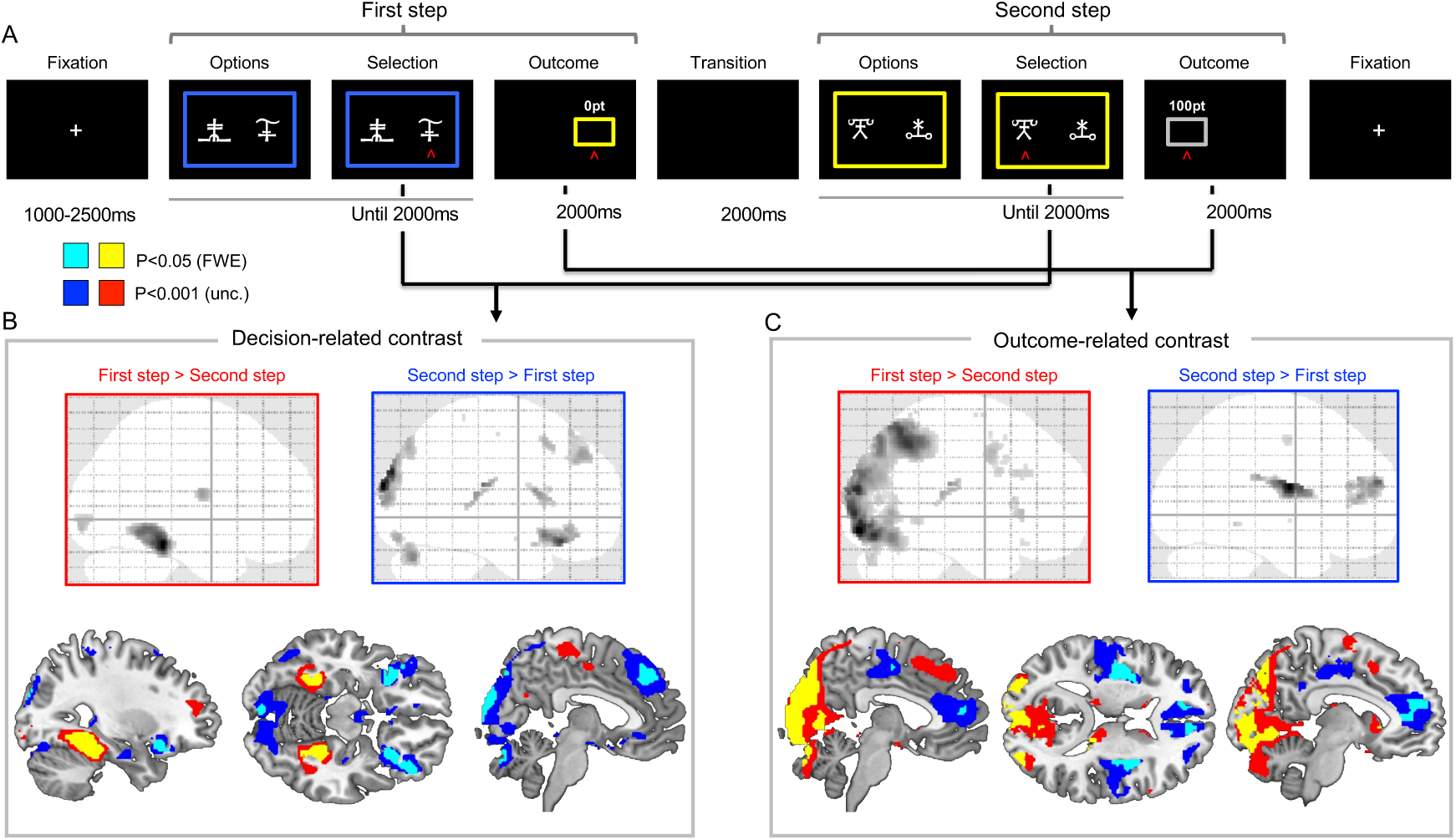
Neural correlates of decision-related contrast and outcome-related contrast. (**A**) Trial organization for learning task: At the start of each trial, a fixation cross was presented for 1000-1200 milliseconds (ms) to introduce the new trial. Two options were then simultaneously presented on the left and right sides, and participants were required to select between them by pressing one of the corresponding buttons until 2000 ms had elapsed. After the selection was made, the outcome appeared for the selected option for 2000 ms before the second step state presentation. The initial decision led to a subsequent state where participants were required to make a new choice. The colored rectangles depict the different states (poor or rich). (**B**) Correlates of brain valuation regions at the moment of choice were examined in a two-step trial onset. Positive activation was observed in the PHC when comparing activations between the first and second steps. Negative activation was observed in the dACC and OFC. (**C**) The study examined the correlates of brain valuation regions during the moment of outcome in a two-step trial onset. Positive activation was observed in the occipital lobe, while negative activation was observed in the insula and mPFC. In (**B**) and (**C**) the voxels in the sagittal glass brain are significant at *P*_*FWE*_ < .05, the activations are also reported on the top of an anatomical template at *P*_*FWE*_ < .05 (yellow and cyan for positive and negative activations, respectively) and *P*_*UNC*_ < .001 (red and blue for positive and negative activations, respectively).

### Forward-looking activations are not dependent on second step state values

While both states of the first step feature the same expected values, this is not true for second step states, because they can be “rich” (average reward ∼50) or “poor” (average reward ∼5). Therefore, GLM1 results (**Fig. 2B** and **2C**), which identified several brain regions differently activated by the first versus the second step (both at the choice and the outcome onset), can be attributed either to the different position of the states in the structure of the task or rather to the differences in their state values. To clarify this issue, we constructed a second GLM (GLM2, **Fig. 3A**) in which trial choice and outcome onsets in the second step were parsed as a function of their state value (“rich” versus “poor”; the structure of GLM2, including parametric modulators, otherwise remained identical to GLM1). We opted for a regions of interest (ROIs) approach and defined our ROIs as the significant clusters identified in GLM1 (*P*_*FWE*_ < .05): PHC, OFC, and dACC (choice-related ROIs); insula, mPFC, occipital and parietal cortices (outcome-related activations).

In the choice-related ROIs (PHC, OFC and dACC), we extracted the regression coefficients of choice onset in the first step and the two second step (“rich” or “poor; **Fig. 3B**). For all three ROIs, regression coefficients of the first step were significantly different from both second step ones (*P* < .001 for all comparisons). Conversely, no significant difference could be detected between the regressors of the two types of second steps (*P* > 0.9, for all comparisons; see **Supplementary Results**). These results suggest that the between-step difference identified in GLM1 was not driven by any particular state (“rich” or “poor”), nor by difference in expected values between them.

We applied a similar approach to the outcome-related ROIs (Occipital, Parietal cortices, mPFC and Insula) and the outcome onsets regression coefficients. For all four ROIs, regression coefficients of the first step were significantly different from both the second step ones (*P* < .001 for all comparisons). Again, the difference between the regressors of the two types of second step did not reach significance (*P* > .081; see **Supplementary Results**). These results suggest that the between-step difference identified in GLM1 was not driven by any particular state (“rich” or “poor”).

**Figure 2.**
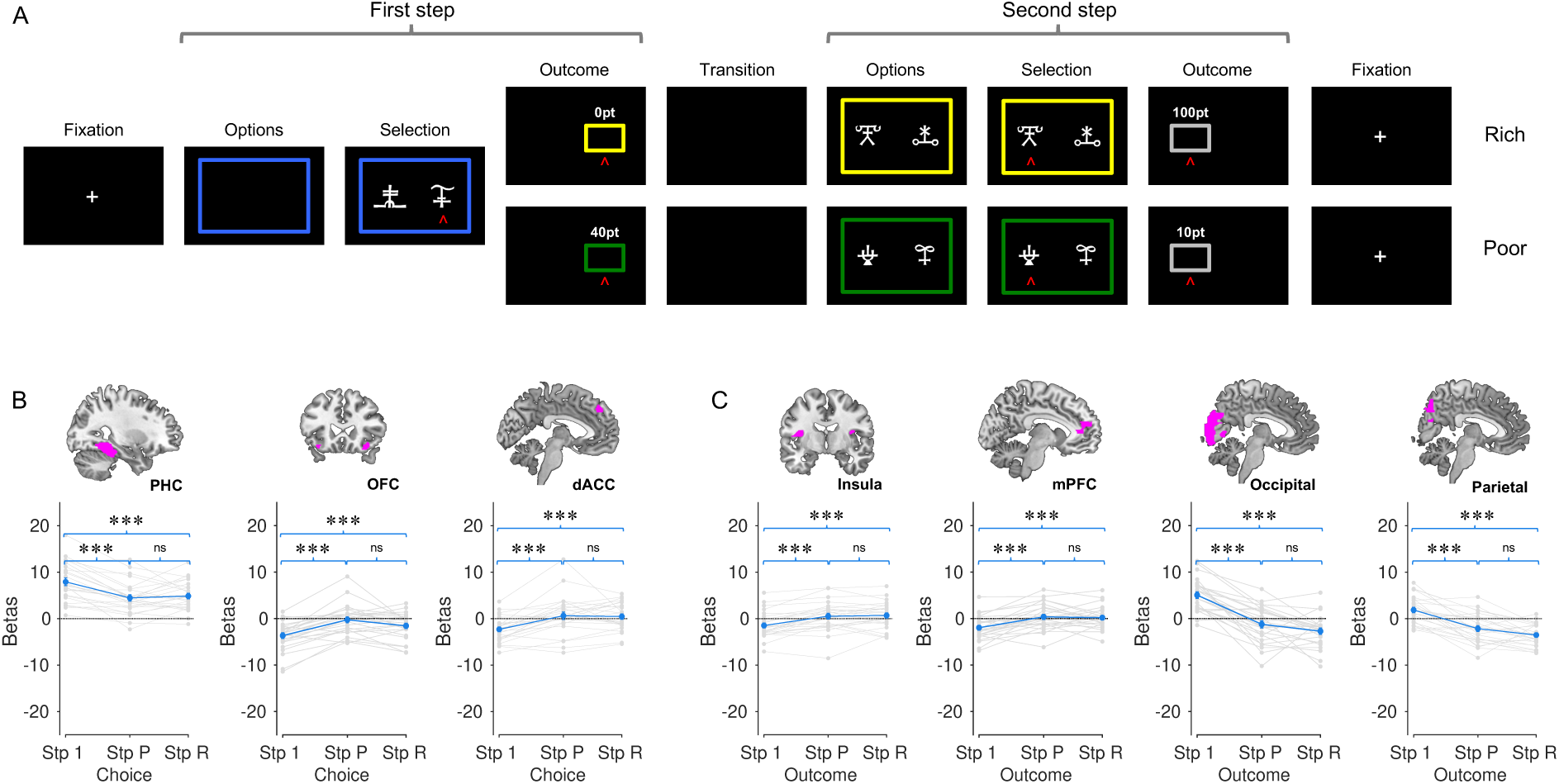
Parametric estimation of the step and state for the second step. **(A)** Parametric estimation was performed on a GLM using a two-step process for trial onset and states in the second step. **(B)** The beta values presented in these graphs were extracted from the ROIs (PHC, OFC, dACC) defined in the first regression, which was based on the decision-related contrast. **(C)** In these graphs, the beta values were extracted from the ROIs defined in the first regression, which was based on the outcome-related contrast. The ROIs include the insula, mPFC, occipital lobe, and parietal cortex. Stp1 (Step 1) corresponds to the onset of the first step; Stp P (Step Poor) corresponds to the onset when participants transition to the poor state in the second step; Stp R (Step Rich) corresponds to the onset when participants transition to the rich state in the second step. The gray lines and dots represent the performance of each participant, while the mean is shown in blue. Error bars indicate the standard error of the mean. Significant differences are denoted by asterisks: * for P < .05, ** for P < .01, and *** for P < .001, Bonferroni corrected one-sample two-sided t-t-test. The significance of the observed condition differences was determined by means of a post-hoc t-test of the one-way repeated measures ANOVA.

### Forward-looking activations as a function of the task-phase

We designed a third GLM (GLM3) to analyze the neural correlates of the progression within the learning sessions (**Fig. 4A**). GLM3 modelled trial onsets (choices and outcomes) separately for trials experienced in the first and the second half of a learning session, which we thereafter refer to the learning-phases. The structure of parametric modulators remained identical to GLM1 and GLM2.

A whole-brain analysis of the learning inception contrast (“first half > second half”) revealed several clusters surviving FWE correction, in the posterior cortex, but also the PHC, striatal and dorsal regions (**Fig. 4A**). The opposite learning progression contrast (“second half > first half”) also revealed smaller activations, mainly concentrated in the insula and the temporal lobe. These activations likely reflect learning-related changes rather than purely anticipatory processes, as they emerge significantly in the contrast comparing early vs. late learning phases. To identify areas responding to both between-step and the between-phase contrast at choice onset, we computed the intersection between the two activations activation maps. This analysis revealed substantial overlap of these two contrasts in the bilateral PHC (*P* < .05, FWE), extending to adjacent areas at more lenient threshold (*P* < .001, uncorrected; **Fig. 4B**). We extracted the betas from ROIs defined in GLM1, which included the PHC, OFC, and the dACC and tested for effects of step and learning-phase, and found that only the PHC displayed a significant effect of learning-phase (see **Fig. S2** and **Supplementary materials**).

We applied the same analyses on outcome-related activations. At the whole brain level, while the learning inception (“first half > second half”) contrast revealed no significant cluster of activation, the opposite learning progression contrast (“second half > first half”) elicited significant activations in large portions of the parietal and frontal regions, as well as in the occipital cortex (*P* < .05, FWE; **Fig. 4C**). We then looked at the intersections of between-step and between-phase activations of the outcome stage. Again, we found substantial overlap in the parietal cortex (*P* < .05, FWE), but also in the dorsal prefrontal cortex and occipital ones, when generating the ROIs at a more lenient threshold (*P* < .001, uncorrected; **Fig. 4D**; see also see **Fig. S3** and **Supplementary materials** for a ROI-based analyses).

In summary, many key brain regions identified in the between-step contrast (namely the PHC at the decision time and the Parietal cortex at outcome onset) had their activity modulated by the progression though the learning sessions. This observation suggests that their activity is dynamically shaped by the learning process.

**Figure 3.**
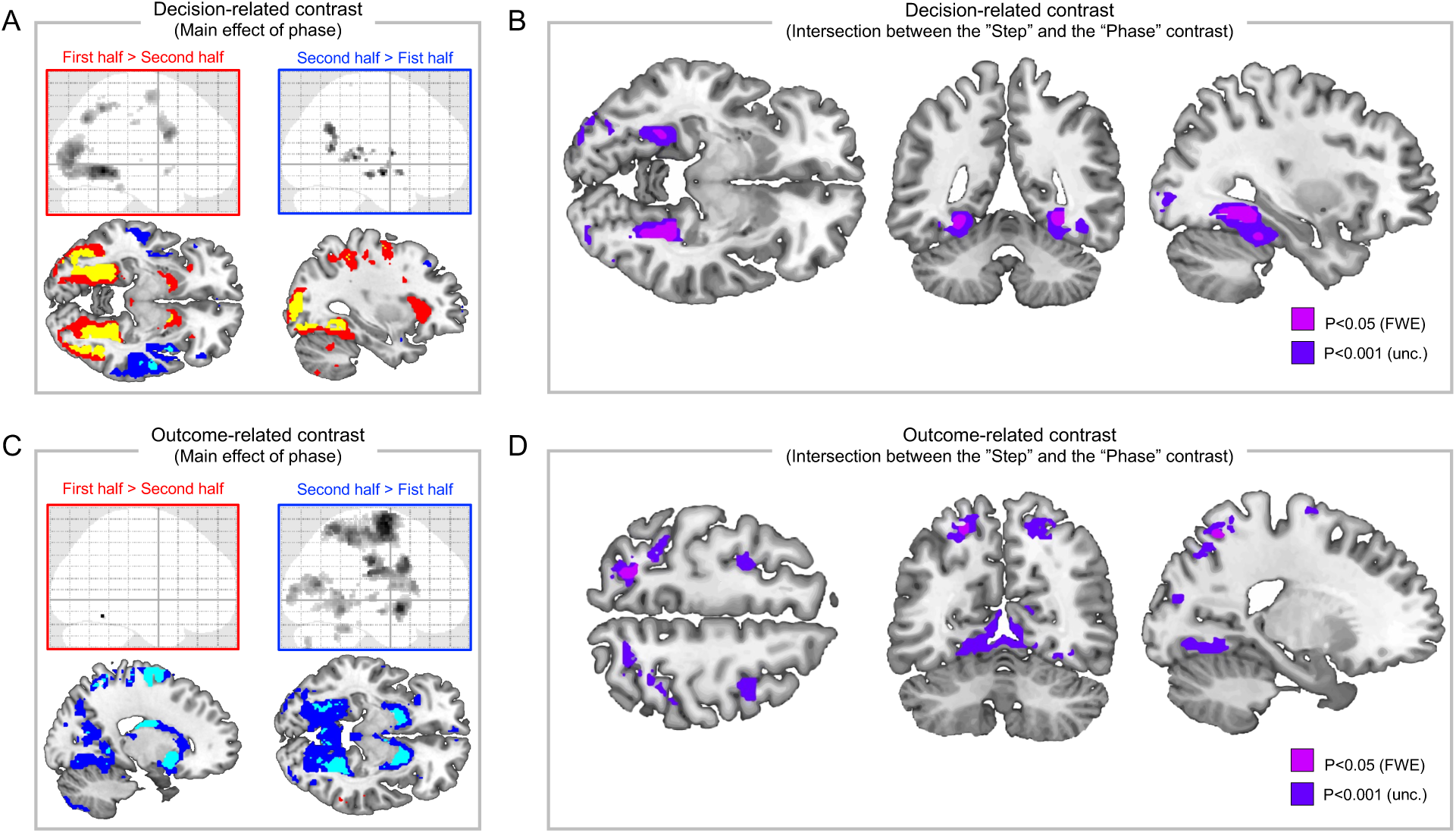
Phase-related contrasts and their overvalue with the step-related contrasts. The figure shows the results of GLM3, where decision and outcome onsets were modelled separately as a function of the task phase (first half: trials 1:40; second half: trials 41:80). (**A**) Phase-related contrast at choice onset. (**B**) Intersection between main effect of step and main effect of phase at the moment of choice. Results indicated significant intersection in the PHC. (**C**) Phase-related contrast at outcome onset. (**D**) Intersection between main effect of step and main effect of phase at the moment of outcome. Results indicated significant intersection in the parietal cortex.

### Neural correlates of reaction times and rewards

In all our GLMs, choice and outcome onsets are respectively parametrically modulated by reaction times ‒ a behavioral proxy for decision-related processing (Clairis & Pessiglione, 2022) ‒ and obtained reward. We therefore assessed the main effect of these regressors, both independently and as a function of the first vs second step (GLM1).

Z-scored reaction times (RTs) were positively correlated with BOLD activity in a frontoparietal network encompassing the insula, corresponding to the saliency and dorsal attentional networks (**Fig. S4**). Notably, RT-related activations remained consistent across steps, with no significant differences between the first and second step at either the whole-brain (*P* > .05, *FWE* − *corrected*) or ROI level. This finding suggests that the neural mechanisms captured by reaction times are therefore unaffected by the necessity of performing forward looking components, as the same frontoparietal and insular regions were consistently engaged in both steps.

At the outcome phase, obtained rewards elicited positive activations in brain regions traditionally associated with the reward system, such as the ventral striatum (VS) and the ventromedial prefrontal cortex (vmPFC), but also in portions of the occipital lobe (*P* < .05, FWE; **Fig. 5A**). These activations appeared stronger for the second than the first step (**Fig. 5B**). To statistically confirm this observation, we extracted the regression coefficients of the reward regressors in ROIs obtained from the activation maps of the step-independent contrast (**Fig. 1A**). In all ROIs, we confirmed that reward encoding significantly increased between the first and second steps (vmPFC: *t*(25) = −3.20, *P* = .004; VS: *t*(25) = −3.53, *P* = .002; Occipital lobe: *t*(25) = −3.34, *P* = .003; **Fig. 5C**).

**Figure 5.**
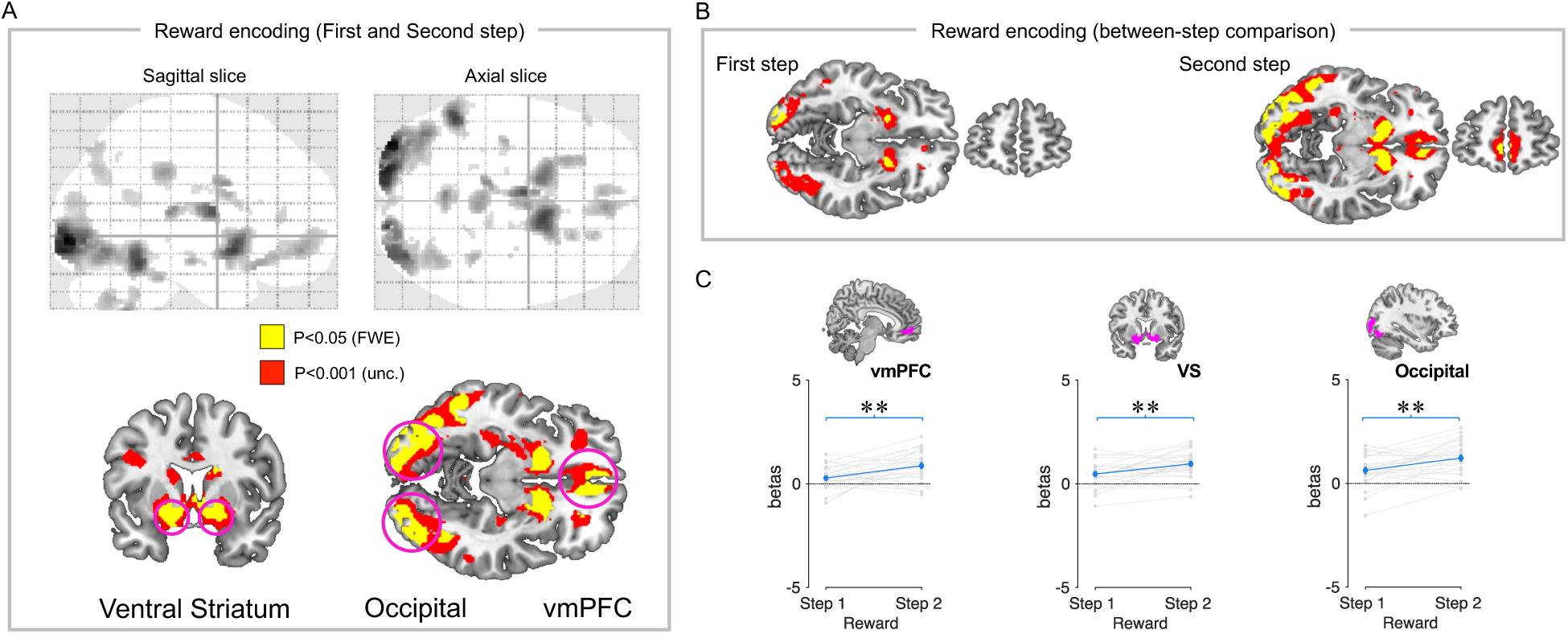
Neural correlates of obtain reward. (**A**) Correlations of brain valuation regions at the moment of outcome for the first step and the second step by the obtained reward parametric modulator. Received reward was positively correlated with activations in the bilateral striatum, occipital lobe, and vmPFC. (**B**) Reward encoding brain activation for the first step and the second step. (**C**) The beta values in these graphs were extracted from the ROIs defined in the first regression, which was based on the parametric reward obtain. The ROIs include the vmPFC, VS, and occipital lobe. The gray lines and dots represent the performance of each participant, while the mean is shown in blue. Error bars indicate the standard error of the mean. Significant differences are denoted by asterisks: * for P < .05, ** for P < .01, and *** for P < .001.

In summary, we replicated traditional results about the neural bases of reaction times and reward encoding in our data. In addition, we show that the multi-step aspect of the task affected reward signals, which were stronger for the second step outcomes.

## Discussion

In this study, we investigated the neural bases of multi-step reinforcement learning in a context where participants had to learn reward values and state transitions simultaneously. To achieve this, we developed a novel variant of the two-step task, where participants navigated through states and made decisions to collect rewards (Daw et al., 2011; Feher da Silva & Hare, 2020; Gläscher et al., 2010; Kool et al., 2016). A distinctive feature of our task is the tension between short-term and long-term rewards implemented at the first-step choices, a common feature in ecological decision-making and inter-temporal choice paradigms in psychology and economics (Frederick et al., 2002; McClure et al., 2004). Additionally, we designed a post-learning assessment to evaluate participants’ explicit knowledge of option values and state transitions after task completion.

Our behavioral results demonstrate that participants successfully learned the task’s contingencies. Second-step choices were consistently above chance levels in both "rich" and "poor" environments, indicating that participants could reliably use learned information to guide decisions. Forward-looking choices at the first step became more frequent in the later phases of the experiment, suggesting a gradual appraisal of the task structure and its progressive inclusions in the decision strategy. Post-learning assessments confirmed that participants accurately retrieved state transitions and identified the values of second-step options. Notably, the reluctance to engage in long-term-oriented decisions during the task was not due to a lack of knowledge of the reward at stakes, since post learning assessments also confirmed that participants were able to correctly identify high values options in the “rich” states. This rather suggest a possible functional dissociation between representing and exploiting a learned model of the task (Hamroun et al., 2022).

To explore the neural mechanisms underlying the strategic tradeoff at stake in our multi-step decision-making task, we first contrasted the BOLD activity between the first-step and second-step decision-making periods, which implement the tension between multi-step planning and single-step reward-maximization. We found that the medial temporal cortex, specifically the parahippocampal cortex was more active during first-step decisions, supporting its role in forward-looking computations and the use of cognitive maps in spatial and conceptual domains (Constantinescu et al., 2016; Garvert et al., 2023). This finding also aligns with prior research linking the medial temporal lobe to future-oriented, imagination-based, decisions in the context of inter-temporal choices (Lebreton et al., 2013; Peters & Büchel, 2010). Interestingly, this activity was higher in the early phases of learning, suggesting that the parahippocampal cortex plays a critical role during the initial stages of forward-planning computations, which potential then transition to other systems as learning progresses and strategies are consolidated. Conversely, the fact that PHC activations were not found at the outcome stage, is in opposition with the idea that hippocampus underpins a replay-based credit-assignment process, which is presumed to be temporally aligned with activity at the final outcome stage of the decision-problem (Liu et al., 2021; Stachenfeld et al., 2017). On the contrary, the PHC seems to encode the distance to the final goal, in a was similar to what has been showed for the hippocampus in human spatial navigation (Howard et al., 2014). Several brain regions such as the dorsomedial prefrontal cortex and orbitofrontal cortex showed an opposite pattern of greater activation during second-step choices, possibly reflecting their involvement in processing immediate rewards and proximity to the task’s endpoint (Farrell et al., 2022).

At outcome presentation stage, we observed a distinct pattern of neural activations. We found that the parietal cortex was more active during first-step outcomes compared to second-step outcomes. Because first-step outcomes combines information about both reward values and state transitions, while second-step outcomes carry information only about reward values, this result is consistent with a role of the parietal cortex in structure learning, which has been documented across various domains (Summerfield et al., 2020; Summerfield & De Lange, 2014; Summerfield & Tsetsos, 2012). Remarkably, the outcome-related parietal activity was modulated by the learning phase, and became more pronounced as learning progressed. While somewhat counter-intuitive, this is in line with the idea that parietal cortex is generally involved in the consolidation of task-relevant knowledge and of action-selection rules, both of which increase with experience (Brodt et al., 2018; Hebscher et al., 2019; Vaidya & Badre, 2022). Altogether, those results support the hypothesis that the parietal cortex is central to the learning and consolidation of task structure, particularly state transitions. The outcome-related contrasts also demonstrate a differential activation pattern between the first and second steps, which can be interpreted through the cognitive processes of credit assignment (Miller & Venditto, 2021; Noonan et al., 2017). Specifically, stronger activations in the medial prefrontal cortex (mPFC) and Insula in the second step may indicate a reassessment of choices and outcomes. This observation is consistent with the concept of credit assignment, where the mPFC and the Insula, may adjust the value of first-step choices based on the received feedback at the second step. These results suggest a complex role of brain networks in outcome processing throughout the task stages.

In contrast with previous studies reporting state prediction errors (SPE) in the dlPFC (Gläscher et al., 2010), our outcome-related contrasts did not reveal an association between activity in the dorsolateral prefrontal cortex (dlPFC) and the processing of state-transition information. However, while these previous studies generally focused on activity associated with varying levels of SPE after extensive training, our study examines the activity elicited by the presence of state-transition-related information (first step) in comparison to its absence (second step). SPEs are commensurate with surprise signals, henceforth can be expected to elicit robust activity in the attentional/saliency network. Consistent with this interpretation, it is worth noting that the same study which found SPE activity in the dlPFC also reported activity related to absolute reward prediction errors (i.e. reward related surprise signals), casting doubts on the specificity of the association between state-transition learning and dlPFC (Fouragnan et al., 2018).

Our results also replicated established findings related to neural representations of reaction times (RTs) and rewards. RTs were associated with widespread activation across the dorsal attentional and saliency networks, independent of task step (Clairis & Pessiglione, 2022; Dixon et al., 2017). Consistent with previous literature reward-related activity was observed in the ventral striatum and prefrontal cortex (Daw et al., 2011; Gläscher et al., 2010; Knutson et al., 2001; McClure et al., 2004). Notably, however present in both step, reward encoding was stronger during second-step outcomes, likely due to the ambiguity surrounding first-step rewards, which depend on participants’ knowledge and utilization of the task model. Families of model-free, model-based and hybrid (e.g., successor-based representations) algorithms propose different ways to compute this potentially multiplexed reward signal, all based on strong assumptions which our design is currently not suited to assess (Daw et al., 2005; Momennejad et al., 2017). A promising approach would be to dynamically modulate the association between outcome types and transitions, in order to orthogonalize reward and structure learning signals.

Our study presents a novel paradigm for investigating the neural mechanisms of multi-step decision-making and model-based computations. We demonstrate a clear functional dissociation between the parahippocampal cortex and the parietal cortex: the former is primarily involved in forward-planning and cognitive mapping, while the latter is critical for structure learning. These findings underscore the dynamic interplay between these regions in orchestrating multi-step decisions, which in turn shape reward-related representations.

Together, this work advances our understanding of the temporal and neural dynamics underlying multi-step reinforcement learning and the associated model-based computations.

## Acknowledgements

SP acknowledges support from the European Research Council consolidator grant (RaReMem: 101043804) three Agence Nationale de la Recherche grants (CogFinAgent: ANR-21-CE23-0002-02; RELATIVE: ANR-21-CE37-750 0008-01; RANGE: ANR-21-CE28-0024-01), the Alexander Von Humbolt foundation and a Google unrestricted gift. ML acknowledges support from the European Research Council Starting grant (INFORL: 948671), and a Google unrestricted gift.

## Methods

### Behavioral task

#### Participants

A total of 57 participants (mean age: 23.06 ± 4.50, 29 females, all right-handed) with no prior history of neurological or psychiatric disorders participated in the experiment. All participants had normal or corrected-to-normal vision and provided informed consent before engaging in the study, which received ethical approval from the relevant authorities. Twenty-eight participants completed the task while inside the fMRI scanner. Two sets of data for participants who underwent the study in a scanner were compromised, resulting in a reduction the number of participants to 26 during the fMRI data analyses.

#### Experimental task

The study comprised of 160 trials, each comprising two steps. Participants were required to select one of two abstract symbols (Agathodaimon alphabet) by pressing a button. At the beginning of each trial, a cross-fixation appeared on the screen. If a participant failed to make a choice within 3 seconds, the task advanced to the next trial. As illustrated in Figure 2, the screen displayed the monetary reward associated with to the color of the subsequent environment. Subsequently, participants selected between symbols in the second step, and the reward was displayed in a neutral (grey) color.

This task was inspired by Glascher et al. (2010), but we chose to provide participants with instructions on both transition structure and associated rewards (see, Table 1). To ensure comprehension, participants were required to pass a quiz before beginning the task. After completing the task, they were asked to assess their understanding of the transition structure (transition assessment post-learning task) and their knowledge of the value associated with each symbol (preference assessment post-learning task). In the transition assessment post-learning task, participants were instructed to estimate the pair of symbols that appeared most frequently following the presentation of a symbol. The presented symbol was one of the four symbols shown in the first step, while the pair of symbols corresponded to one of the two pairs from the second step.

### fMRI recording and analysis

#### fMRI data recording

The participants underwent scanning with a Siemens 3T Tim-Trio MRI scanner, using a 32-channel surface coil. Acquisition parameters included TE = 25.0 ms and TR = 1022.0 ms. The scans covered the entire brain with slices of 2.5 mm thickness and a slice-gap of 3.00. The voxel size was 2.5 x 2.5 x 2.5 mm3, and the echo spacing was 0.5 ms. Additionally, a whole-brain high-resolution structural T1-weighted MPRAGE sequence was performed to characterize participants’ anatomy, with slices having a voxel size of 1 mm^3^ and acquisition parameters of TR = 2300 ms and TE = 4.18 ms.

#### fMRI pre-processing

FSL was utilized, and standard preprocessing procedures were applied (Jenkinson et al., 2012). These included EPI realignment and unwrapping using field maps, segmenting T1 images into gray matter, white matter, and cerebrospinal fluid, and using segmentation parameters to warp T1 images to the SPM Montreal Neurological Institute (MNI) template. The normalized functional data was spatially smoothed using an isotropic 8-mm full-width half-maximum Gaussian kernel. For each participant, a mask was applied to the whole brain anatomical gray matter, derived by averaging the normalized T1 anatomical scans of our participants and later thresholding the image at 0.5.

#### fMRI data analysis

The fMRI data were analyzed using a General Linear Model (GLM) framework, implemented in the Statistical Parametric Mapping version 12 (SPM12) package (Ashburner et al., 2014). The results presented in this study are derived from three related GLMs. The statistical threshold for the main results was set at family-wise error (FWE - Bonferroni) correction *P* < .05. For illustrative purposes, finding is also presented using an uncorrected threshold of *P* < .001.

The initial GLM1 was conceived with the objective of identifying distinct cerebral regions that exhibited differential activation during the comparison of the first and second steps in our multi-step decision-making learning task. The model incorporated four categorical regressors, denoted as indicator functions, to capture the onset of: (1) choice at the first step; (2) outcome presentation at the first step; (3) choice at the second step; and (4) outcome presentation at the second step. Additionally, four parametric regressors were incorporated to account for: (1) reaction time at the first step; (2) reward obtained at the first step; (3) reaction time at the second step; and (4) reward obtained at the second step. Prior to their inclusion in the analysis, the reaction time regressors were standardized (z-scored) (Lebreton et al., 2019).

The aim of GLM2 was to ascertain whether significant differences between steps identified in GLM1 were not driven by comparison with specific step 2 states (‘rich’ or ‘poor’). The model incorporated six categorical regressors, modeled as indicator functions denoting the onset of: (1) choice at the first step, (2) outcome presentation at the first step, (3) choice at the second step in the poor environment, (4) outcome presentation at the second step in the poor environment, (5) choice at the second step in the rich environment, and (6) outcome presentation at the second step in the rich environment. Additionally, six parametric regressors were incorporated to account for: (1) reaction time at the first step, (2) reward obtained at the first step, (3) reaction time at the second step in the poor environment, (4) reward obtained at the second step in the poor environment, (5) reaction time at the second step in the rich environment, and (6) reward obtained at the second step in the rich environment. All parametric regressors for reaction time underwent standardization (z-scored).

A last GLM (GLM3) was used to investigate the neural correlates associated with progression across trials within each learning session. For each participant, the trials were divided into two blocks: the first 40 trials (‘First Half’) and the last 40 trials (‘Second Half’). Eight categorical regressors were included as indicator functions to model the onset of: (1) choice at the first step in the first half, (2) outcome presentation at the first step in the first half, (3) choice at the first step in the second half, (4) outcome presentation at the first step in the second half, (5) choice at the second step in the first half, (6) outcome presentation at the second step in the first half, (7) choice at the second step in the second half, and (8) outcome presentation at the second step in the second half. Four parametric regressors were also included to account for: (1) reaction time at the initial stage, (2) reward obtained at the initial stage, (3) reaction time at the subsequent stage, and (4) reward obtained at the subsequent stage. All reaction time regressors were standardized (z-scored).

#### ROI data-analysis

Subsequent to analysis in the GLM1, we identified a set of regions activated within the significant clusters, corrected for familywise error (*P*_*FWE*_ < .05). At the choice onset significant between step activations were observed in the parahippocampal cortex (PHC), orbitofrontal cortex (OFC), and dorsal anterior cingulate cortex (dACC). At the outcome onset, we identified significant activations in the insula, medial prefrontal cortex (mPFC), occipital lobe, and parietal cortex when contrasting the outcome onsets at the first step with those at the second step. Beta values were extracted for all ROIs in the GLM2 and the GLM3. A one-way repeated-measures ANOVA was performed for GLM2 and two-way repeated-measures ANOVA for the GLM3, followed by post-hoc tests with Bonferroni correction, to compare beta values across the different categories.

For the "reward encoding" contrast in GLM1, reward-related activations were observed in the ventromedial prefrontal cortex (vmPFC), ventral striatum (VS), and occipital lobe. Beta values for these three ROIs were extracted, and comparisons of beta values were performed between Step 1 and Step 2.

## Supplementary Materials

### Supplementary Results

#### GLM2: detailed statistical comparisons

Here we provide the details concerning the statistical tests performed on the ROIs defined based on GLM1 and the regression coefficients extracted from GLM2, in order to assess that the between-step results obtained in GLM1 where not driven by differences with specific a second step states (“rich” or “poor”) or, in other terms, were truly induced by the structure of the task and not by the value of the states.

Concerning choice-related activations and ROIs, significant differences were observed at the moment of choice between the initial and subsequent steps (either poor or rich) across all examined ROIs. The PHC demonstrated heightened brain activity during the first step compared to the rich state in the second step (*t*(25) = 5.97, *P* < .001) and the poor state in the second step (*t*(25) = 6.49, *P* < .001). In contrast, both the OFC and the dACC exhibited decreased activity in the first step compared to the rich state (OFC: *t*(25) = −5.14, *P* < .001, dACC: *t*(25) = −5.31, *P* < .001) and poor state (OFC: *t*(25) = −6.65, *P* < .001, dACC: *t*(25) = −5.08, *P* < .001) in the second step. No significant differences in brain activity were observed between the poor and rich steps (PHC: *t*(25) = −0.68, *P* = 1, OFC: *t*(25) = 2.31, *P* = .089, dACC: *t*(25) = 0.27, *P* = 1). This observation indicates that the results of the initial general linear model (GLM1) were not influenced by the state conditions during the decision-making process.

Concerning outcome related activations and choices, it was observed that there were significant differences between the first and second steps in both the rich and poor states. However, no significant differences were found between the rich and poor states at the second step across all ROIs. In particular, insula and mPFC exhibited decreased brain activity during the first step in comparison to the rich state (insula: *t*(25) = −5.34, *P* < .001, mPFC: *t*(25) = −4.53, *P* < .001) and poor state (insula: *t*(25) = −6.00, *P* < .001, mPFC: *t*(25) = −4.67, *P* < .001) in the second step. In contrast, the occipital lobe and the parietal cortex showed increased brain activity between the first step compared to rich state (Occipital: *t*(25) = 9.62, *P* < .001, Parietal: *t*(25) = 9.11, *P* < .001) and poor state (Occipital: *t*(25) = 7.85, *P* < .001, Parietal: *t*(25) = 6.07, *P* < .001) in the second step. No significant differences in brain activity were observed between the poor and rich states (insula: *t*(25) = −0.44, *P* = 1, mPFC: *t*(25) = 0.35, *P* = 1, occipital: *t*(25) = 1.79, *P* = .256, and parietal: *t*(25) = 2.35, *P* = .081).

To sum up, there ample positive statistical evidence that the between-step (first step, versus second step) comparisons where not driven by difference to a particular second step state (“rich” or “poor”). This conclusion is supported by the significant results of all comparisons between the regression coefficient of the first step and those of the second step. Additionally, there is not positive statistical evidence of any difference between “rich” and “poor” states in any of the specified ROIs and contrasts.

#### GLM3: detailed statistical comparisons

In this section, we detail the statistical tests conducted on the ROIs defined from GLM1 and the regression coefficients extracted from GLM3. The objective of these analyses was to verify that the between-step results obtained in GLM1 were driven by differences between the first 40 trials (First Half) and the last 40 trials (Second Half), but rather reflected the progression across the learning process.

Concerning choice-related activations and ROIs, significant differences were observed at the moment of choice between first half and second half trials in the two steps (step 1 and step 2) for the PHC but not for the OFC and the dlPFC ROIs. In PHC brain activity decreased between first half and second half for the step 1 (*t*(25) = 4.78, *P* < .001) and the step 2 (*t*(25) = 8.36, *P* < .001). Conversely, no such differences were observed in the first step for the OFC (*t*(25) = −1.79, *P* = .512) and the dlPFC (*t*(25) = −2.72, *P* = .07). Similarly, no differences were observed in step 2 for the OFC (*t*(25) = .531, *P* = 1) and the dlPFC (*t*(25) = .29, *P* = 1).

Concerning outcome-related activations, it was observed that there were significant differences in the first step between the first half and the second half trials for the Insula (*t*(25) = −3.55, *P* = .009) but not in the second step (*t*(25) = −2.48, *P* = .119). No significant differences where observed between first half and second half trials in step 1 for the mPFC (*t*(25) = .005, *P* = 1), occipital lobe (*t*(25) = −1.07 = 1) and the Parietal (*t*(25) = −2.09, *P* = .281, nor in the second step (mPFC: *t*(25) = .12, *P* = .909; occipital: *t*(25) = −2.53, *P* = .109; Parietal: *t*(25) = −2.10, *P* .276).

In summary, the significant changes observed are primarily located in the PHC for choice-related activations and in the insula for outcome-related activations, highlighting specific evolutions throughout the learning process. This observation suggests that their role is not merely statistical but is actively shaped by the dynamics of the learning process.

---

**Figure S1.**
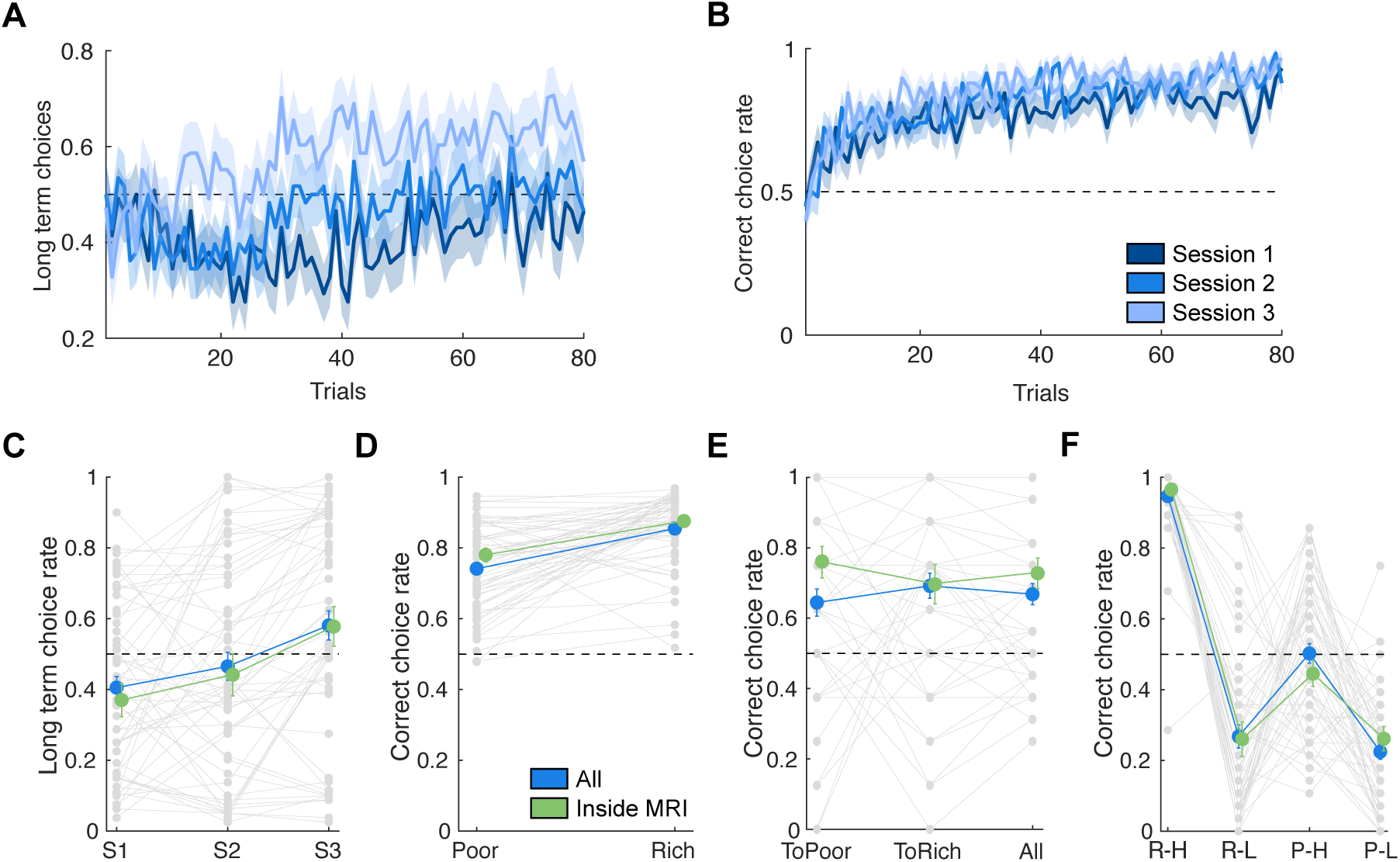
Behavioral results for all participants (blue) and MRI population (green). (**A**) Time course of long-term choices (choosing the option in the first step has a high probability of going to the rich state in the second step) for the 3 sessions. (**B**) Time course of correct choice rate (chosen the best option) in the second step for the three sessions (**C**) The graph represents the long-term choice rate in the learning task per sessions. (**D**) The graph represents the rate of correct choices based on the state in the second step (Poor or Rich). (**E**) The graph displays the rate at which participants accurately selected the first step option that led to either the poor or rich state in the second step. (**F**) The graph displays the accuracy rate at the second step based on the state (poor/rich) and outcome value (low/high). As illustrated in panels A and B, the color gradient is indicative of the respective session colors. The darkest shade corresponds to session 1, while the lightest shade corresponds to session 3. The lines and dots in panels C, D, E, and F in blue correspond to the results for all participants, and in green to the results for participants who completed the study in the MRI. The gray lines and dots represent the performance of each participant, while the mean is shown in blue for all participants (included behavioral and fMRI population) and fMRI population. Error bars indicate the standard error of the mean. Significant differences are denoted by asterisks: * for *P* < .05, ** for *P* < .01, and *** for *P* < .001.

**Figure S2.**
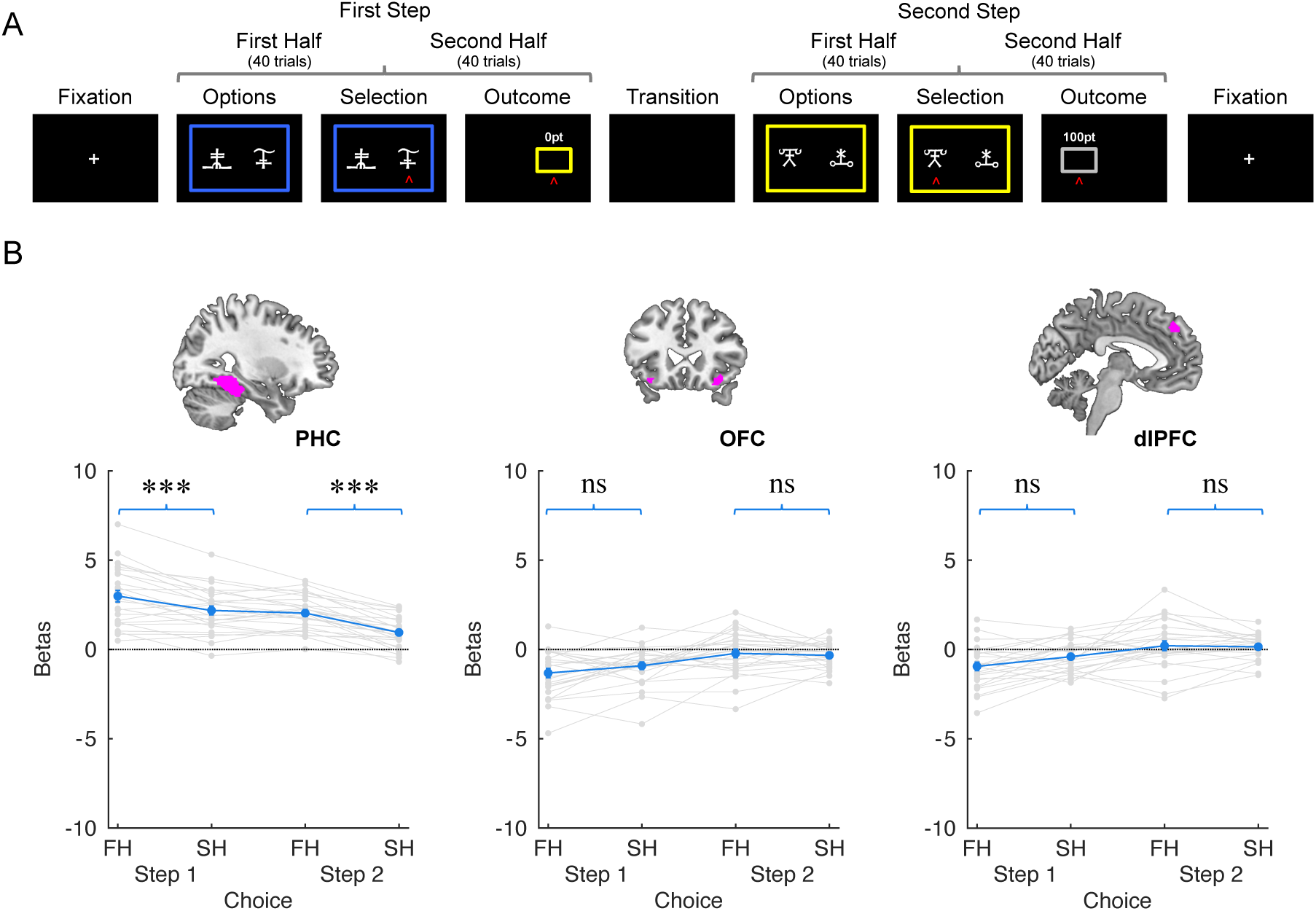
Parametric estimation of two steps and progression across trials within each learning session in choice onset. (**A**) Parametric estimation was performed on a GLM using a two-step process (step 1 and step 2) and trials split half trials (first half 1:40 and second half 41:80) (**B**) The beta values presented in these graphs were extracted from the ROIs (PHC, OFC and dlPFC) defined in the first regression, which was based on the decision-related contrast. The gray lines and dots represent the performance of each participant, while the mean is shown in blue. Error bars indicate the standard error of the mean. Significant differences are denoted by asterisks: * for *P* < .05, ** for *P* < .01, and *** for *P* < .001, A two-tailed, one-sample t-test with Tukey correction was used. The significance of the observed differences between conditions was assessed using a post-hoc t-test following a one-way repeated measures ANOVA.

**Figure S3.**
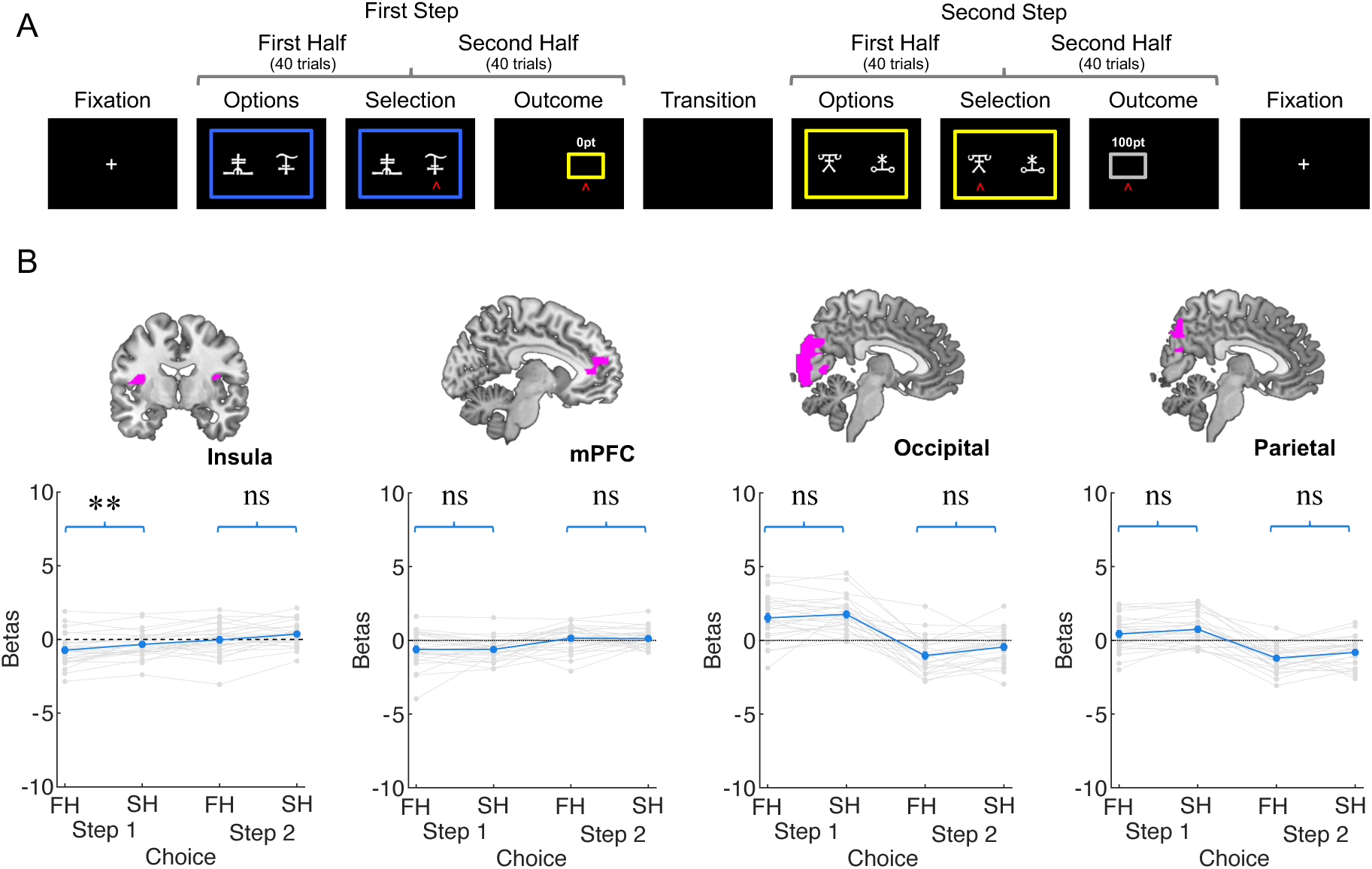
Parametric estimation of two steps and progression across trials within each learning session in outcome onset. (**A**) Parametric estimation was performed on a GLM using a two-step process (step 1 and step 2) and trials split half trials (first half 1:40 and second half 41:80) (**B**) The beta values presented in these graphs were extracted from the ROIs (insula, mPFC, occipital lobe and Parietal cortex) defined in the first regression, which was based on the outcome-related contrast. The gray lines and dots represent the performance of each participant, while the mean is shown in blue. Error bars indicate the standard error of the mean. Significant differences are denoted by asterisks: * for *P* < .05, ** for *P* < .01, and *** for *P* < .001, A two-tailed, one-sample t-test with Bonferroni correction was used. The significance of the observed differences between conditions was assessed using a post-hoc t-test following a one-way repeated measures ANOVA.

**Figure S4.**
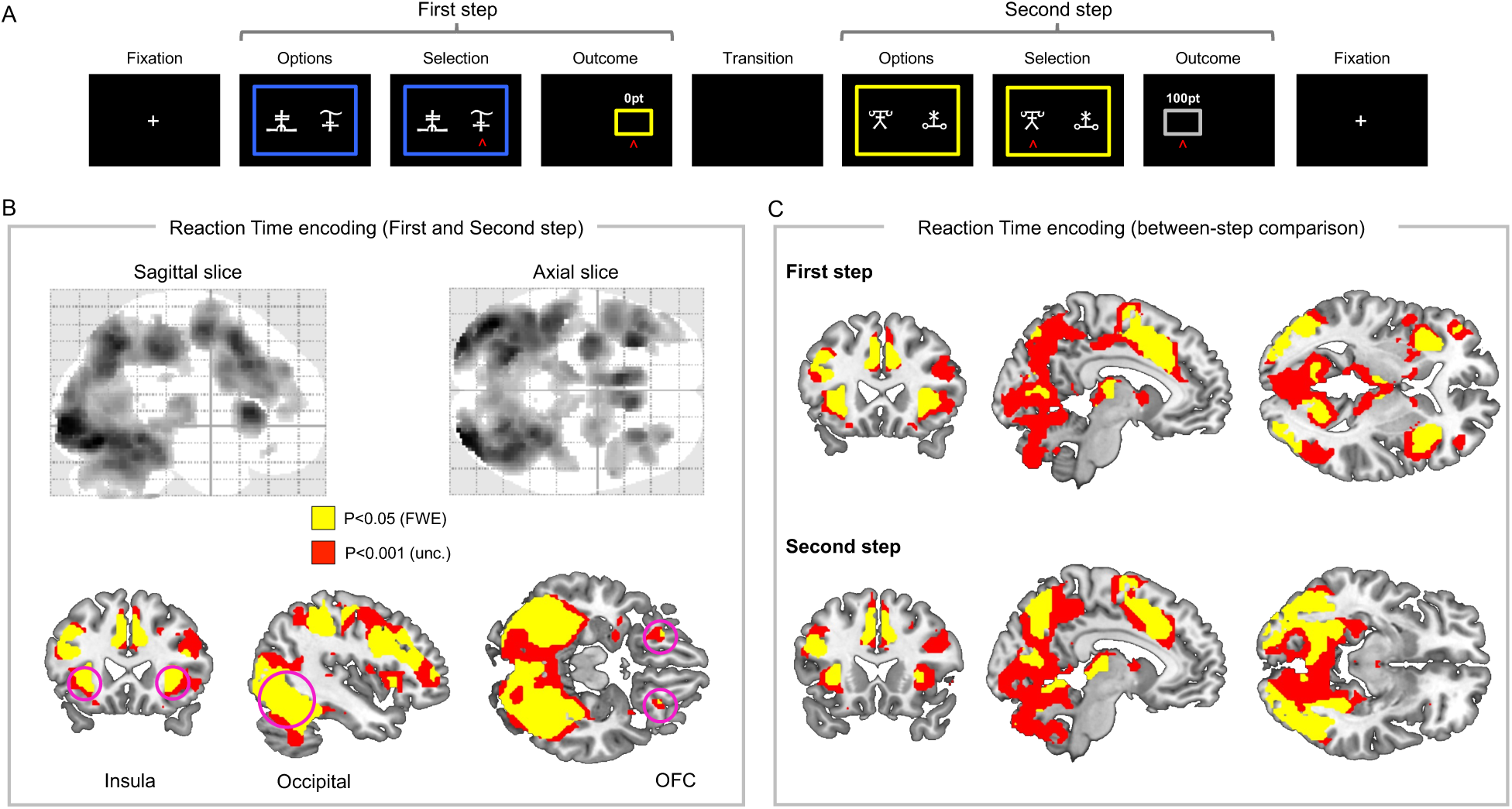
Neural correlates of reaction time. (**A**) Trial organization for the learning task (**B**) Correlations of brain valuation regions at the moment of choice for the first step and the second step of the reaction time parametric modulator. Reaction time was positively correlated with activations in the insula, occipital lobe, and OFC. (**C**) Reaction time encoding brain activation for the first step and the second step independently. In both the first step and the second step of the experiment, reaction time was found to be positively correlated with activations in the insula, hippocampus, ACC, and occipital lobe.

**Table S1.**
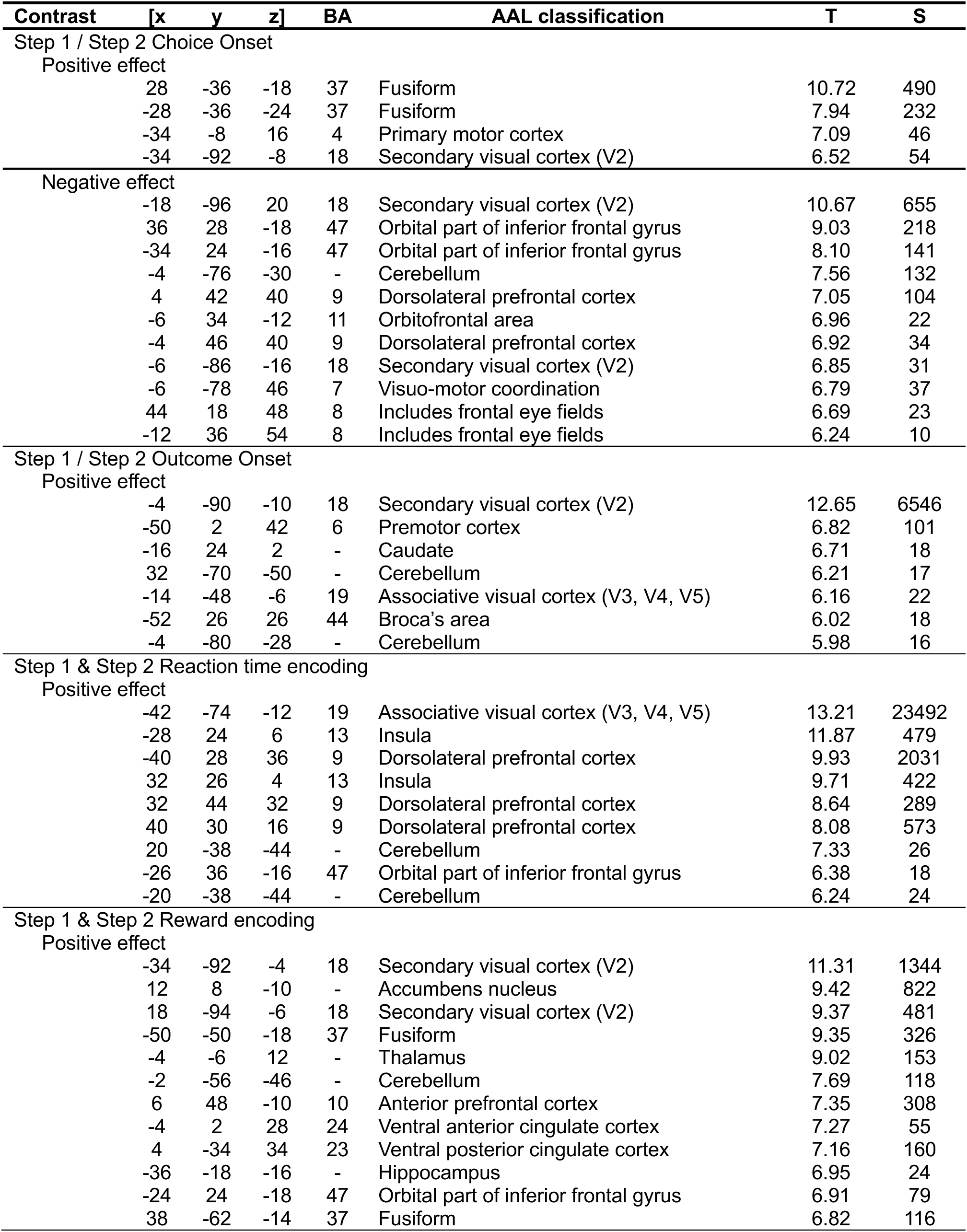
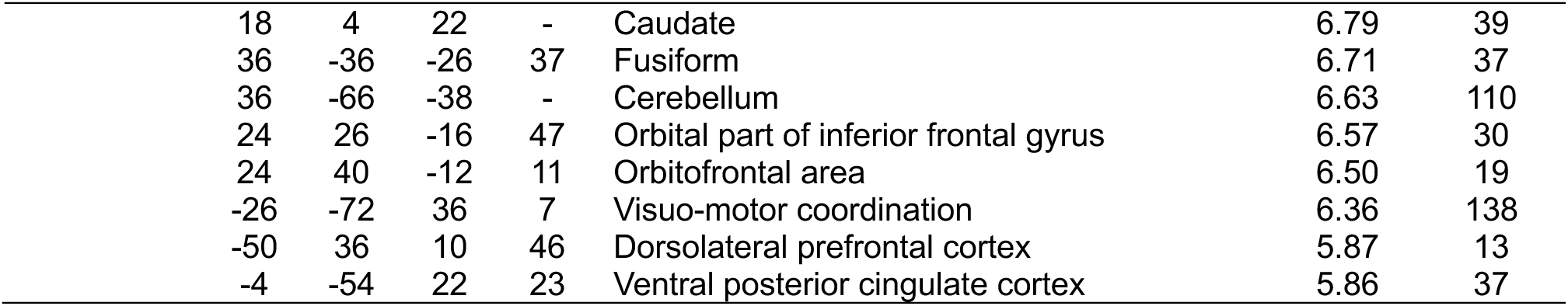
Decision-related contrast, outcome-related contrast between step 1 and step 2 (GLM1), reaction time encoding and rewards encoding ROIs. The following table contains a list of activated brain areas with a number of voxels in the cluster superior at 10. [x, y, z] correspond to the coordinates of activation pic in MNI space. BA refers to Brodmann Area, a cortical region defined by its cytoarchitecture and associated with specific brain functions. AAL refers to the Automated Anatomical Labeling atlas (Tzourio-Mazoyer et al., 2002), a parcellation system of the brain used for functional and structural neuroimaging analyses. T refers to the T-value at the activation peak within the cluster, indicating the statistical significance of the observed effect, and S refers to the number of voxels in the cluster.

| Contrast | [x | y | z] | BA | AAL classification | T | S |
| --- | --- | --- | --- | --- | --- | --- | --- |
| Step 1 / Step 2 Choice Onset |  |  |  |  |  |  |  |
| Positive effect |  |  |  |  |  |  |  |
|  | 28 | -36 | -18 | 37 | Fusiform | 10.72 | 490 |
|  | -28 | -36 | -24 | 37 | Fusiform | 7.94 | 232 |
|  | -34 | -8 | 16 | 4 | Primary motor cortex | 7.09 | 46 |
|  | -34 | -92 | -8 | 18 | Secondary visual cortex (V2) | 6.52 | 54 |
| Negative effect |  |  |  |  |  |  |  |
|  | -18 | -96 | 20 | 18 | Secondary visual cortex (V2) | 10.67 | 655 |
|  | 36 | 28 | -18 | 47 | Orbital part of inferior frontal gyrus | 9.03 | 218 |
|  | -34 | 24 | -16 | 47 | Orbital part of inferior frontal gyrus | 8.10 | 141 |
|  | -4 | -76 | -30 | - | Cerebellum | 7.56 | 132 |
|  | 4 | 42 | 40 | 9 | Dorsolateral prefrontal cortex | 7.05 | 104 |
|  | -6 | 34 | -12 | 11 | Orbitofrontal area | 6.96 | 22 |
|  | -4 | 46 | 40 | 9 | Dorsolateral prefrontal cortex | 6.92 | 34 |
|  | -6 | -86 | -16 | 18 | Secondary visual cortex (V2) | 6.85 | 31 |
|  | -6 | -78 | 46 | 7 | Visuo-motor coordination | 6.79 | 37 |
|  | 44 | 18 | 48 | 8 | Includes frontal eye fields | 6.69 | 23 |
|  | -12 | 36 | 54 | 8 | Includes frontal eye fields | 6.24 | 10 |
| Step 1 / Step 2 Outcome Onset |  |  |  |  |  |  |  |
| Positive effect |  |  |  |  |  |  |  |
|  | -4 | -90 | -10 | 18 | Secondary visual cortex (V2) | 12.65 | 6546 |
|  | -50 | 2 | 42 | 6 | Premotor cortex | 6.82 | 101 |
|  | -16 | 24 | 2 | - | Caudate | 6.71 | 18 |
|  | 32 | -70 | -50 | - | Cerebellum | 6.21 | 17 |
|  | -14 | -48 | -6 | 19 | Associative visual cortex (V3, V4, V5) | 6.16 | 22 |
|  | -52 | 26 | 26 | 44 | Broca's area | 6.02 | 18 |
|  | -4 | -80 | -28 | - | Cerebellum | 5.98 | 16 |
| Step 1 & Step 2 Reaction time encoding |  |  |  |  |  |  |  |
| Positive effect |  |  |  |  |  |  |  |
|  | -42 | -74 | -12 | 19 | Associative visual cortex (V3, V4, V5) | 13.21 | 23492 |
|  | -28 | 24 | 6 | 13 | Insula | 11.87 | 479 |
|  | -40 | 28 | 36 | 9 | Dorsolateral prefrontal cortex | 9.93 | 2031 |
|  | 32 | 26 | 4 | 13 | Insula | 9.71 | 422 |
|  | 32 | 44 | 32 | 9 | Dorsolateral prefrontal cortex | 8.64 | 289 |
|  | 40 | 30 | 16 | 9 | Dorsolateral prefrontal cortex | 8.08 | 573 |
|  | 20 | -38 | -44 | - | Cerebellum | 7.33 | 26 |
|  | -26 | 36 | -16 | 47 | Orbital part of inferior frontal gyrus | 6.38 | 18 |
|  | -20 | -38 | -44 | - | Cerebellum | 6.24 | 24 |
| Step 1 & Step 2 Reward encoding |  |  |  |  |  |  |  |
| Positive effect |  |  |  |  |  |  |  |
|  | -34 | -92 | -4 | 18 | Secondary visual cortex (V2) | 11.31 | 1344 |
|  | 12 | 8 | -10 | - | Accumbens nucleus | 9.42 | 822 |
|  | 18 | -94 | -6 | 18 | Secondary visual cortex (V2) | 9.37 | 481 |
|  | -50 | -50 | -18 | 37 | Fusiform | 9.35 | 326 |
|  | -4 | -6 | 12 | - | Thalamus | 9.02 | 153 |
|  | -2 | -56 | -46 | - | Cerebellum | 7.69 | 118 |
|  | 6 | 48 | -10 | 10 | Anterior prefrontal cortex | 7.35 | 308 |
|  | -4 | 2 | 28 | 24 | Ventral anterior cingulate cortex | 7.27 | 55 |
|  | 4 | -34 | 34 | 23 | Ventral posterior cingulate cortex | 7.16 | 160 |
|  | -36 | -18 | -16 | - | Hippocampus | 6.95 | 24 |
|  | -24 | 24 | -18 | 47 | Orbital part of inferior frontal gyrus | 6.91 | 79 |
|  | 38 | -62 | -14 | 37 | Fusiform | 6.82 | 116 |

|  |  |  |  |  |  |  |
| --- | --- | --- | --- | --- | --- | --- |
| 18 | 4 | 22 | - | Caudate | 6.79 | 39 |
| 36 | -36 | -26 | 37 | Fusiform | 6.71 | 37 |
| 36 | -66 | -38 | - | Cerebellum | 6.63 | 110 |
| 24 | 26 | -16 | 47 | Orbital part of inferior frontal gyrus | 6.57 | 30 |
| 24 | 40 | -12 | 11 | Orbitofrontal area | 6.50 | 19 |
| -26 | -72 | 36 | 7 | Visuo-motor coordination | 6.36 | 138 |
| -50 | 36 | 10 | 46 | Dorsolateral prefrontal cortex | 5.87 | 13 |
| -4 | -54 | 22 | 23 | Ventral posterior cingulate cortex | 5.86 | 37 |

## Notes

### Competing Interest Statement

The authors have declared no competing interest.

